# Hyperspectral ocean color encodes global eukaryotic phytoplankton community structure

**DOI:** 10.64898/2026.09.09.750502

**Authors:** Sasha J. Kramer, Roy El Hourany

## Abstract

Phytoplankton community composition (PCC) impacts global ocean change across scales. Combining the taxonomic resolution of in situ methods with the spatiotemporal coverage of satellite ocean color remains a major challenge. Phytoplankton pigments link PCC to ocean color; past work has demonstrated that 4-6 pigment-based groups can be reconstructed from hyperspectral remote sensing reflectance (*R_rs_*(*λ*)). Alternately, metabarcoding approaches reveal thousands of taxa to the species level, but the links between optics and genes are less clear. Here, we compiled a global, coincident dataset of hyperspectral *R_rs_*(*λ*), pigments, and 18S rRNA gene sequences. We used a bio-optical model to produce the relative contributions of 13 eukaryotic phytoplankton classes identified by metabarcoding with high confidence in most cases. Model coefficients were highly spectrally correlated between genes and pigments, revealing the strong PCC signal within *R_rs_*(*λ*). This proof-of-concept study establishes a foundation for detecting eukaryotic PCC beyond pigments in the PACE era.

## Introduction

Phytoplankton community composition (PCC) and biodiversity exert first-order control on a wide array of marine ecological and biogeochemical processes, including the biological carbon pump (Le Quéré et al. 2005; Guidi et al. 2009), oceanic nutrient cycling (Arrigo 2005), ecosystem functioning and resilience (Baert et al. 2016; Behrenfeld et al. 2021), and the marine food webs that support pelagic fisheries (Legendre 1990). As a result, characterizing spatiotemporal dynamics in surface ocean PCC is essential to describe biological functioning in the global ocean from the cell to ecosystem level (de Vargas et al. 2015; Righetti et al. 2019; Sommeria-Klein et al. 2021). However, a major challenge remains in balancing the high taxonomic resolution provided by in situ approaches with the broad spatiotemporal coverage of ocean color satellites, which currently resolve comparatively coarse biological properties (Cetinić et al. 2024). Bridging this observational gap requires determining how much taxonomically-resolved PCC information is encoded in ocean color and can ultimately be recovered at the scales that shape marine ecosystems in a changing ocean.

Phytoplankton pigments provide the primary link between PCC and ocean color remote sensing reflectance (*R_rs_*(*λ*)). Pigment data support the robust statistical separation of 4-7 broad chemotaxonomic groups depending on the dataset (Kramer and Siegel 2019), and impart a discernible signal in ocean color data due to changes in light absorption based on the composition and concentration of pigments in those groups (Bracher et al. 2017; Mouw et al. 2017; Werdell et al. 2018). However, pigment-based PCC represents only one dimension of community composition since pigments can be shared among phytoplankton taxa and their relative concentrations can vary with physiological state (Jeffrey et al. 2011). NASA’s Plankton Aerosol Cloud ocean Ecosystem (PACE) satellite and its Ocean Color Imager (OCI) offer the first global, hyperspectral view of the surface ocean and therefore an opportunity to separate fine-scale spectral features associated with PCC beyond chlorophyll-*a* (Werdell et al. 2019, 2026; Cetinić et al. 2024). PACE’s hyperspectral resolution supports the development of approaches that target fine-scale optical features and correlate those features with phytoplankton properties, including (but not limited to) pigment concentration and composition (Wolanin et al. 2016; Vandermeulen et al. 2020).

Three PCC algorithms have thus far been selected for implementation with PACE. The Multi-Ordination ANAlysis (MOANA) model derives *Prochlorococcus*, *Synechococcus*, and picoeukaryote cell abundances from hyperspectral *R_rs_*(*λ*) (Lange et al. 2020), while the Gaussian Pigments (GPig) and Spectral Derivative Pigments (SDP) algorithms retrieve phytoplankton pigments and associated broad taxonomic groups from their optical signatures (Chase et al. 2017; Kramer et al. 2022). Together, these approaches demonstrate that hyperspectral *R_rs_*(*λ*) contains biologically meaningful information beyond chlorophyll-*a*, but their taxonomic resolution remains constrained to the information contained in the flow cytometry or pigment data used to develop the models (Kramer et al. 2024a).

Molecular approaches, in contrast, can describe PCC at substantially finer taxonomic resolution and have increasingly revealed relationships between plankton community structure, ecosystem functioning, and biogeochemical processes (e.g., (Guidi et al. 2016; Ibarbalz et al. 2019; Stephens et al. 2024; Kramer et al. 2025; Lampe et al. 2025; Kramer 2026). Historically, ocean color remote sensing approaches have been siloed from genetic methods and these fields have developed largely independently. A growing number of studies have started to merge these methods (Kaneko et al. 2023; El Hourany et al. 2024; Marchese et al. 2026), but only for multispectral data. For the next generation of PACE algorithms, optics and genes must be combined to resolve the PCC information content of ocean color data and create models that go *beyond pigments* to assess the potential and limitations of retrieving advanced phytoplankton taxonomy from space.

To address this knowledge gap, here we combine in situ measurements of PCC from 18S rRNA gene metabarcoding with in situ hyperspectral *R_rs_*(*λ*) data to test whether molecularly-defined eukaryotic PCC leaves a recoverable optical imprint in ocean color data. Using 40 paired observations, we compare the PCC derived independently from pigments and metabarcoding data to evaluate how successfully each representation can be reconstructed from hyperspectral *R_rs_*(*λ*). Specifically, we ask whether hyperspectral *R_rs_*(*λ*) contains information associated with eukaryotic PCC assessed from metabarcoding, test whether this community composition can be reconstructed with meaningful predictive skill, and assess whether the recoverable molecular PCC signal extends beyond the information represented by the pigment-based groups separated in this dataset. The paired observations show a high dynamic range in PCC and optical conditions, providing a proof-of-concept that will motivate future comparisons with broader taxonomic range (e.g., including prokaryotic gene markers) and encourage future field campaigns to collect paired measurements in more and different global ecosystems.

## Methods

### Compiling a global dataset

Publicly-available datasets were compiled based on the following criteria: samples were collected in the surface ocean (<=10 meters) and included HPLC phytoplankton pigment data collected following best practices, 18S rRNA gene sequence data, and hyperspectral remote sensing reflectance (*R_rs_*(*λ*)), all collected at the same times and places following the criteria in (Kramer et al. 2024a), for a total of 40 paired surface samples (**Figure 1A**). Samples were included from: the North Atlantic Aerosols and Marine Ecosystems Study (NAAMES; N = 6) in the western North Atlantic Ocean, collected across seasons in 2015 to 2018 (Behrenfeld et al. 2019); EXport Processes in the Ocean from RemoTe Sensing (EXPORTS) in the North Pacific (August-September 2018; N = 4) and North Atlantic (May 2021; N = 15) Oceans (Siegel et al. 2021; Johnson et al. 2023); and the Tara Oceans field campaign (N = 15) from 2009 to 2012 (Sunagawa et al. 2020).

**Figure 1.**
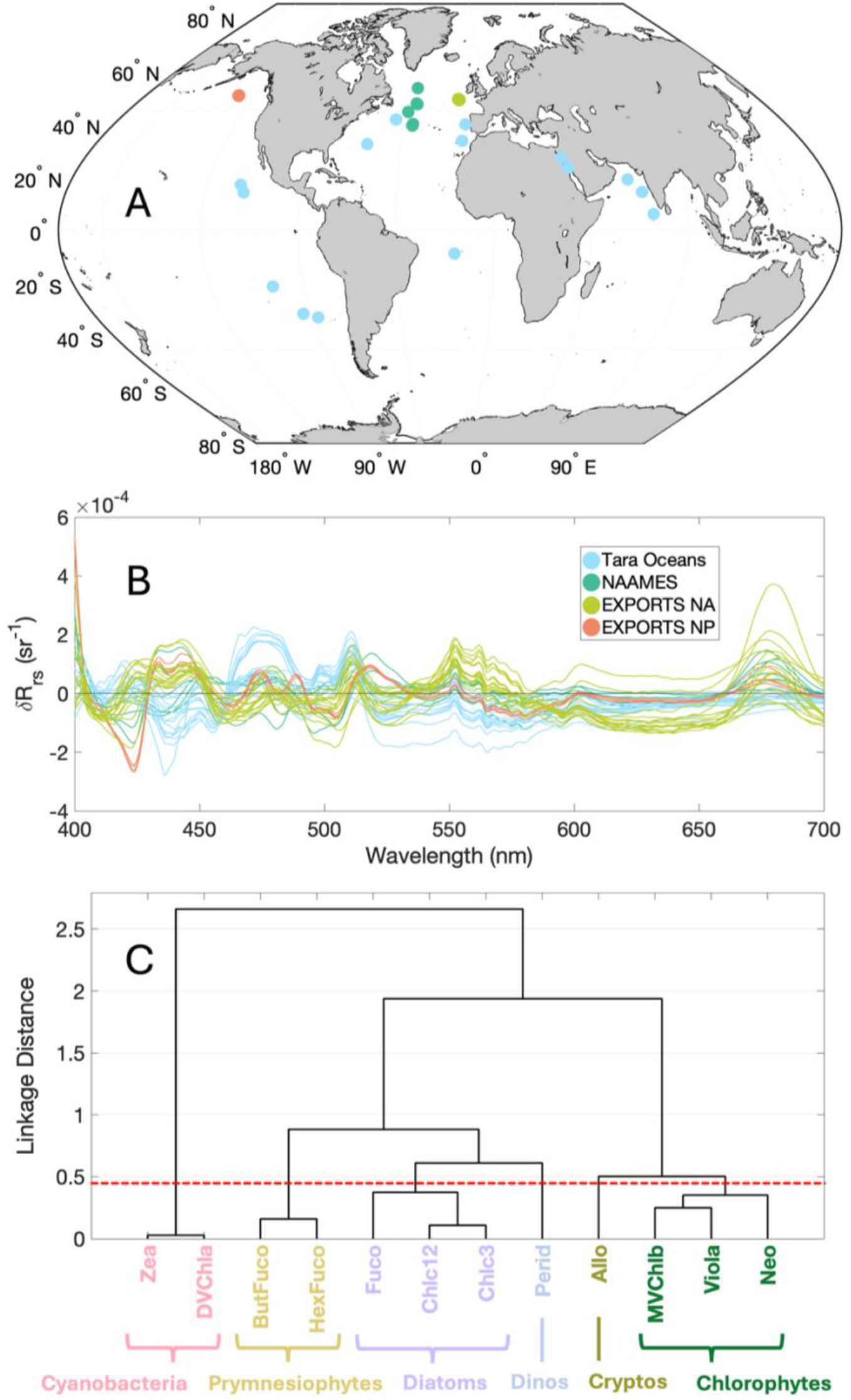
(A) Global map of paired hyperspectral *R_rs_*(*λ*), HPLC, and 18S rRNA gene sequences used in this analysis (blue = Tara Oceans, dark green = NAAMES, olive green = EXPORTS NA, orange = EXPORTS NP). (B) SDP model-generated *δR_rs_*(*λ*) spectra for all data (colored the same as in A). (C) Pigment-based phytoplankton groups separated by hierarchical cluster analysis in this dataset: cyanobacteria, prymnesiophytes, diatoms, dinoflagellates, cryptophytes, chlorophytes.

### HPLC pigments

Each field campaign measured concentrations of total chlorophyll-*a* (Tchla) and phytoplankton accessory pigments. Consistent pigments were measured at each sampling site, but the twelve accessory pigments that are retrieved by SDP (Kramer et al. 2022) were the focus in this study: peridinin (Perid), fucoxanthin (Fuco), chlorophyll c1+c2 (Chlc12), chlorophyll c3 (Chlc3), hexanoyloxyfucoxanthin (HexFuco), butanoyloxyfucoxanthin (ButFuco), alloxanthin (Allo), prasinoxanthin (Pras), neoxanthin (Neo), monovinyl chlorophyll b (MVchlb), divinyl chlorophyll a (DVchla), and zeaxanthin (Zea). This subset of accessory pigments was selected for their taxonomic relevance for describing photosynthetic phytoplankton community composition (Kramer and Siegel 2019; Kramer et al. 2022). A hierarchical cluster analysis was performed on the relative concentration of phytoplankton pigments to Tchla using the correlation distance and Ward’s linkage method (**Figure 1C**). Six distinct chemotaxonomic phytoplankton groups separated from this analysis: dinoflagellates, diatoms, prymnesiophytes, chlorophytes, cryptophytes, and cyanobacteria (Jeffrey et al. 2011; Kramer and Siegel 2019). The pigment ButFuco was also used to compare to silicoflagellate abundance (pelagophytes and dictyochophytes) from metabarcoding data. Total chlorophyll-*a* concentrations in the global dataset ranged from 0.02-1.28 mg m^-3^ with a mean concentration of 0.48 mg m^-3^.

### 18S rRNA gene sequences

Eukaryotic phytoplankton community composition was characterized using 18S rRNA gene sequences. The 18S rRNA gene is limited to describing only eukaryotic phytoplankton groups, so the prokaryotic fraction of the phytoplankton community is not assessed here. However, the 18S rRNA gene was selected over the 16S rRNA gene for this study to capture variability in relative dinoflagellate sequence abundances, as this group cannot fully be quantified or described using 16S rRNA genes (Lin 2011). For most broad taxonomic groups, relative 18S rRNA gene sequence abundance has been shown to covary with relative phytoplankton biomass measured across methods that quantify cell counts and/or cellular carbon biomass (Lin et al. 2019; Catlett et al. 2020, 2022; Kramer et al. 2024a). Relative sequence abundance was therefore used here to assess relative PCC rather than as a direct estimate of cell abundance. However, since the 18S rRNA gene is not found in cyanobacteria, the sequence data used here do not represent prokaryotic phytoplankton (captured by variability in HPLC pigments measured from the same samples, e.g., relative contributions by Zea and DVchla).

The specific methods for gene amplification and sequencing for all datasets used in this study have been described in detail elsewhere and therefore are briefly summarized here. The V4 region of the 18S rRNA gene was sequenced for Tara Oceans (de Vargas et al. 2015; Delage et al. 2023) and EXPORTS (SeaBASS 2018; Kramer et al. 2025); amplicon sequence variants (ASVs) were inferred using DADA2 and taxonomically assigned against PR2 v5.0.0 (Guillou et al. 2013; Callahan et al. 2016). In the NAAMES dataset, the V9 region of the 18S rRNA gene was sequenced and taxonomy was assigned using PR2 (SeaBASS 2014; Kramer et al. 2024a). Phytoplankton taxonomy has been found to be comparable at the class level between 18S rRNA V4 and V9 (Zimmermann et al. 2024). After assigning taxonomy, ASVs were classified as either photosynthetic or heterotrophic in accordance with prior trophic classifications (Durkin et al. 2022; Kramer et al. 2025). ASVs assigned to known heterotrophic dinoflagellates and parasitic lineages (e.g., *Syndiniales* spp.; (Guillou et al. 2008)) were removed (Jones et al. 2025). Remaining dinoflagellate ASVs were assumed to be photosynthetic, though some dinoflagellates (and other phytoplankton, including many prymnesiophytes) are facultatively mixotrophic. The resulting dataset therefore retained ASVs classified as photosynthetic or potentially photosynthetic.

Photosynthetic phytoplankton ASVs were summed into 13 shared taxonomic classes for consistency across datasets: Pelagophyceae and Dictyochophyceae (silicoflagellates); Chrysophyceae; Bolidophyceae; Bacillariophyceae (diatoms); Chlorarachniophyceae (chlorophyll b-containing); Prymnesiophyceae (prymnesiophytes); Cryptophyceae (cryptophytes); Mamiellophyceae, Pyramimonadophyceae, and other Chlorophyceae (chlorophytes); Dinophyceae (dinoflagellates); and other Ochrophyta (red algae).

### Hyperspectral remote sensing reflectance (R_rs_ (λ))

Finally, hyperspectral remote sensing reflectance was measured at each sampling site. All *R_rs_*(*λ*) spectra used in this analysis were also used in (Kramer et al. 2024b) and additional details on the acquisition and processing of the *R_rs_*(*λ*) spectra used here can be found in (Chase et al. 2017). The measured spectral range varied among field campaigns; therefore, only values from 400-700 nm were used for consistency. All spectra were interpolated to a common 1 nm wavelength resolution and smoothed using a 5 nm moving mean filter before further analysis.

### Principal components regression modeling

As described above, the Spectral Derivative Pigments (SDP) algorithm (Kramer et al. 2022) was developed to model phytoplankton pigment concentrations from hyperspectral *R_rs_*(*λ*). Full details of the SDP model are found in (Kramer et al. 2022) and model code is publicly available on GitHub (see *Data availability statement*). Briefly, the model constructs a residual spectrum (*δR_rs_*(*λ*); **Figure 1B**) representing the difference between a measured *R_rs_*(*λ*) spectrum and a modeled *R_rs_*(*λ*) spectrum based on a generic optical model. The SDP model is then trained using a principal components regression (PCR) approach to retrieve coefficients and intercepts for each pigment at each wavelength, which are used to reconstruct the pigment concentrations. Following the original SDP implementation, model performance was evaluated using 100 repeated random train-validation partitions. At each cross-validation, 75% of the samples were randomly assigned to model training and the remaining 25% were withheld for validation. Repeating this procedure 100 times allowed the sensitivity of model performance to the train-validation partition to be assessed for this relatively small dataset. In the original SDP model, the second derivative of *δR_rs_*(*λ*), *δR_rs_*"(*λ*), was used to model the concentrations of total chlorophyll-*a* and the 12 accessory pigments described above. Here, we retrained the SDP model using *δR_rs_*"(*λ*) to reconstruct the concentrations of these 13 pigments, and then applied the same PCR framework separately to reconstruct the relative contributions of the 13 phytoplankton classes assessed from 18S rRNA gene sequences.

## Results and Discussion

### Modeling HPLC pigment concentrations

The concentrations of all 13 pigments were reconstructed with strong agreement across the dataset and within individual samples (**Figure 2**). The coefficient of determination between measured and modeled pigments, *R*^2^, ranged from 0.65 to 0.93 (**Figure S1**). The performance of the retrained SDP model on this dataset exceeded the performance of the original SDP model for many accessory pigments (Kramer et al. 2022), demonstrating the strong relationship between *δR_rs_*(*λ*) features and phytoplankton pigment concentrations in this subset of that dataset. Importantly, the relationships between and among relative accessory pigment concentrations were also maintained across the dataset, with the same number of phytoplankton chemotaxonomic groups separating from a hierarchical cluster analysis of the modeled pigments as separated from the measured pigments (Figure 1C; (Kramer and Siegel 2019)). Both dataset-wide patterns and much of the sample-level variability were reproduced by the model (**Figure 2**), supporting the ability of the *δR_rs_*(*λ*) method to capture broad and subtle patterns in spectral variability, associated with phytoplankton pigment composition and concentration in this dataset (Kramer et al. 2024b). This result emphasizes the theoretical basis of the SDP model for pigments, which aims to remove broad-scale spectral variability associated with absorption and scattering by non-phytoplankton parameters, maximizing the relationships between phytoplankton taxonomy, pigments, absorption, and reflectance (Siegel et al. 2005, 2013; Catlett and Siegel 2018).

**Figure 2.**
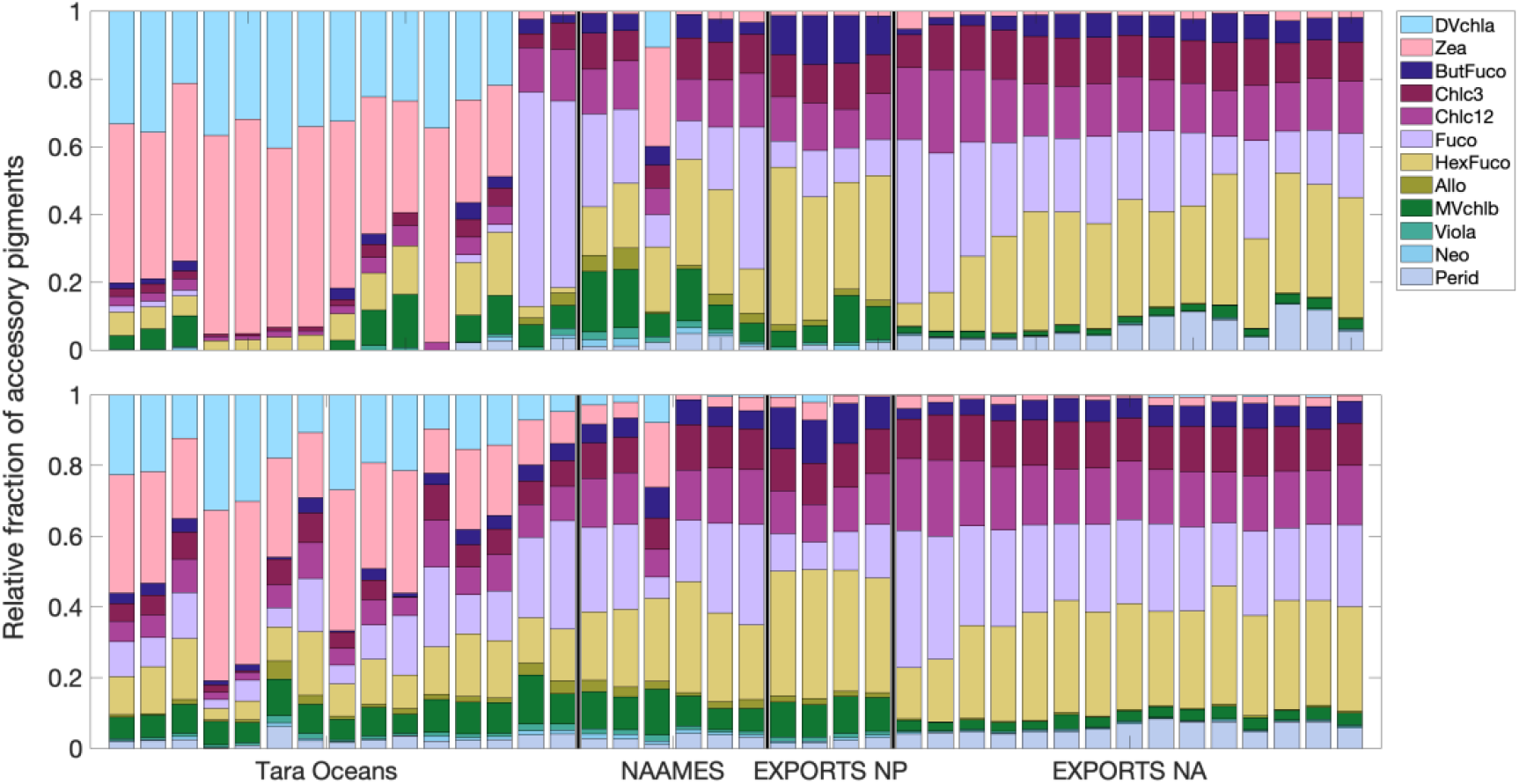
Measured (upper) and *δR_rs_*(*λ*)-modeled (lower) relative fractions of phytoplankton accessory pigments for all samples in Figure 1.

### Modeling 18S rRNA gene relative sequence abundances

While phytoplankton pigments are expected to have a strong optical signal associated with both taxonomy and ocean color (Bidigare et al. 1989; Ciotti et al. 1999; Bricaud et al. 2004), it is less certain if phytoplankton genetic variability will impart a recoverable signal in *R_rs_*(*λ*). Here, eukaryotic PCC derived from 18S rRNA gene sequences showed substantial correspondence with the information contained in *δR_rs_*(*λ*). The 18S rRNA gene models reproduced the variability across all 13 phytoplankton classes (**Figure 3**), with *R*^2^ values ranging from 0.53 to 0.79 (**Figure S2**).

**Figure 3.**
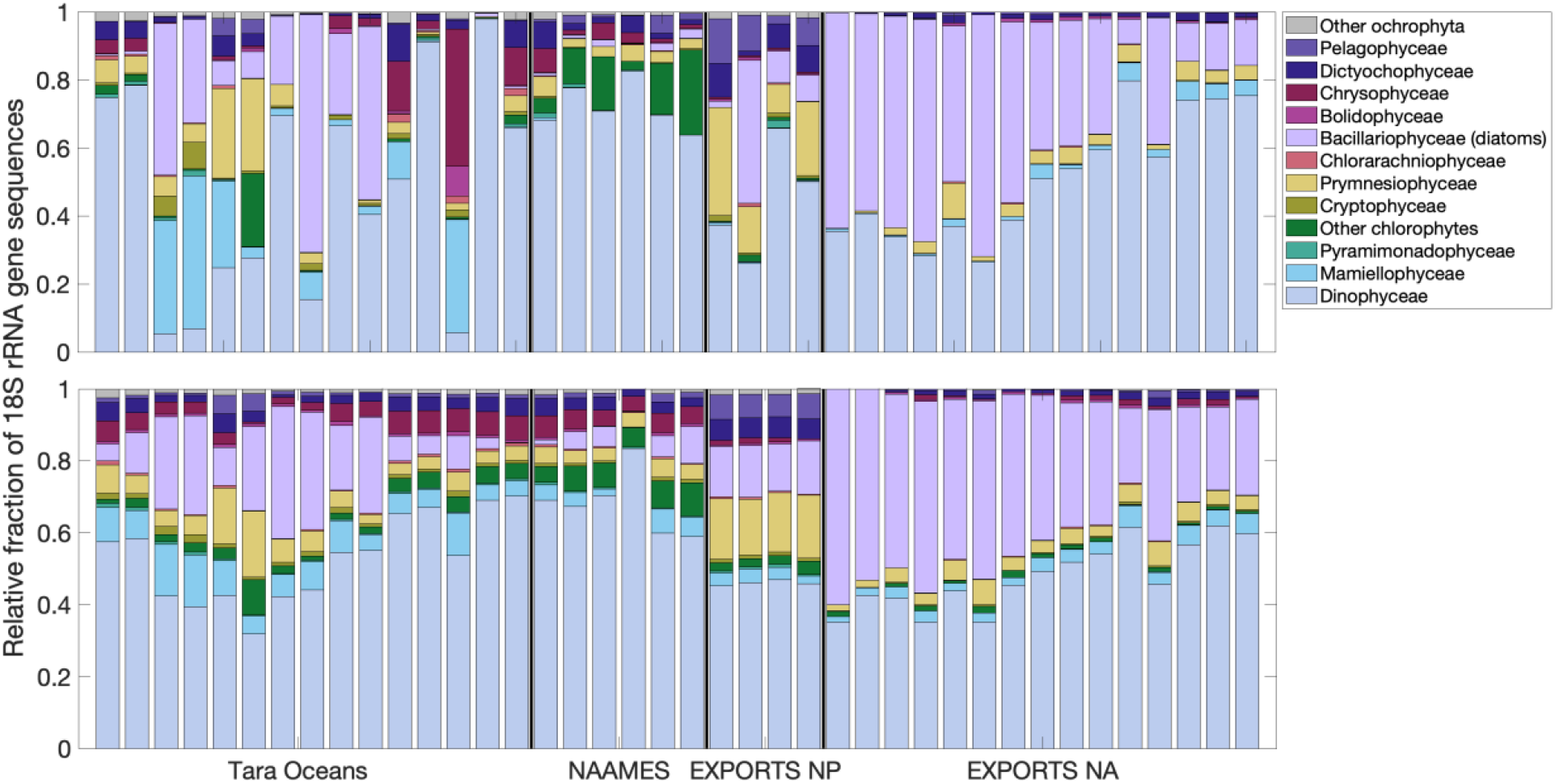
Measured (upper) and *δR_rs_*(*λ*)-modeled (lower) relative fractions of 18S rRNA gene phytoplankton classes for all samples in Figure 1.

As with the pigment models, the 18S rRNA gene models reconstructed the broad PCC variation across the dataset, particularly capturing broad trends between diatoms and dinoflagellates. However, sample-level differences were reproduced less consistently than for pigments, indicating that some molecular PCC variation was not represented by the available optical information. In a first example, during the EXPORTS North Pacific field campaign, the optical signal showed relatively little variation across 4 separate sampling days (**Figure 1B**) that captured phytoplankton communities with similar pigment composition (**Figure 2**) but different phytoplankton composition from metabarcoding (**Figure 3**). The resulting modeled 18S rRNA gene PCC therefore did not capture the variability observed between samples, since the only input to the model was the *δR_rs_*(*λ*) spectrum. More fundamentally, this result emphasizes that the taxonomic resolution available from molecular observations does not necessarily translate directly into optically-recoverable signal from ocean color. 18S rRNA gene-based PCC is expected to be distinguishable from hyperspectral *R_rs_*(*λ*) when changes in community composition are accompanied by corresponding changes in the integrated optical properties of the water column. Conversely, taxonomic turnover that produces little change in pigment composition (as seen in **Figure 2**), cell-level optical properties, or other sources of spectral variability may remain effectively invisible to an optical retrieval.

In a second example, the missing photosynthetic prokaryotic community composition can help explain some of the confusion in the eukaryotic-only 18S rRNA gene model: several samples measured on Tara Oceans with, e.g., high relative chrysophyte or chlorophyte sequence abundance, were reconstructed without those contributions (**Figure 3**). In the pigment data from those samples (**Figure 2**), strong relative contributions from pigments associated with prokaryotic phytoplankton (namely DVchla, found only in *Prochlorococcus* spp.) indicate that substantial components of the optically-active phytoplankton community were absent from the 18S rRNA gene-derived target composition (Jeffrey et al. 2011). Thus, an ocean color signal that integrates both eukaryotic and prokaryotic cell information cannot necessarily map uniquely onto a PCC target that represents only the eukaryotic fraction.

### Eukaryotic PCC from pigments and genes

Despite differences between pigment- and metabarcoding-derived PCC in this dataset (**Figures 2-3**), the relative contributions of several major eukaryotic groups were positively related between the two approaches (**Figure S3**). Spearman’s correlation coefficients (*ρ*) for diatoms and cryptophytes were particularly strong (0.92 and 0.91, respectively), while *ρ* values for prymnesiophytes and silicoflagellates were slightly lower (0.48 and 0.21, respectively) though still significant (p < 0.05). The relationships between pigments and genes for these groups have been shown to vary across other ecosystems and communities, likely due to differences in feeding strategy and pigment composition within broadly diverse taxa (Jordan and Chamberlain 1997; Zapata et al. 2004; Catlett et al. 2022; Kramer et al. 2024a). Additionally, the 18S rRNA gene relative sequence abundance does not map directly onto either cell abundance or biomass because rRNA gene copy numbers vary substantially among taxa, particularly for dinoflagellates. The major difference in PCC between these two methods for this dataset is the inclusion of prokaryotic phytoplankton in the pigment data, which are absent from the 18S rRNA dataset. The signature of the cyanobacterial pigments is strong across samples collected on Tara and NAAMES (**Figure 2**), and prediction performance for some eukaryotic groups was lower within subsets of samples from these field campaigns (**Figure S2**). Since cyanobacteria (particularly *Prochlorococcus* spp.) are known to be globally important contributors to marine ecosystems (Follows et al. 2007; Biller et al. 2015; Li et al. 2022), future optical-molecular comparisons should incorporate markers quantifying both prokaryotes and eukaryotes to represent the full photosynthetic phytoplankton community and to better separate taxonomic variation from mismatches in biological coverage among methods.

### Spectral correlations between optics, pigments, and genes

The relationships observed in this dataset among optics, pigments, and genes revealed wavelength regions that were consistently associated with particular taxonomic groups across pigment- and gene-based models (**Figure 4**). Specifically, the models for diatoms, prymnesiophytes, and cryptophytes showed similar spectral regions associated with the corresponding accessory pigment- and 18S rRNA gene-derived PCC signals. This correspondence suggests that some of the molecular community composition signal covaries with optical features that are also associated with pigment composition, providing a mechanistic bridge between taxonomic composition and hyperspectral ocean color. However, there are also examples where the spectral signals for pigments and genes are very divergent (for instance, dinoflagellates) or vary between 18S rRNA gene classes and bulk accessory pigments (chlorophytes). Dinoflagellates are well known for high variability in cell size and shape, feeding strategy, pigment content (or lack thereof), and gene copies—characterizing such divergent taxa as members of a broader taxonomic group likely leads to differences in the optical vs. genetic signals, as are seen here (Lin 2011; Zapata et al. 2012; Lin et al. 2019). The consistent spectral signature associated with cyanobacterial pigments in this dataset identifies wavelength regions that could be tested explicitly against prokaryotic PCC markers in future studies (e.g., using 16S rRNA genes). The results shown here also compare well to relationships between optics, pigments, and genes in a separate global analysis using a different gene marker (El Hourany and Kramer 2026), encouraging future studies that directly assess the phytoplankton gene-based information content in *R_rs_*(*λ*). Additionally, while the current analysis was limited by in situ matchup data, PACE is now in space collecting daily hyperspectral *R_rs_*(*λ*). PACE data opens the door for further analyses that use ocean color information collected concurrently with genetic information, even from studies where dedicated in situ hyperspectral *R_rs_*(*λ*) measurements were not able to be made in the field.

**Figure 4.**
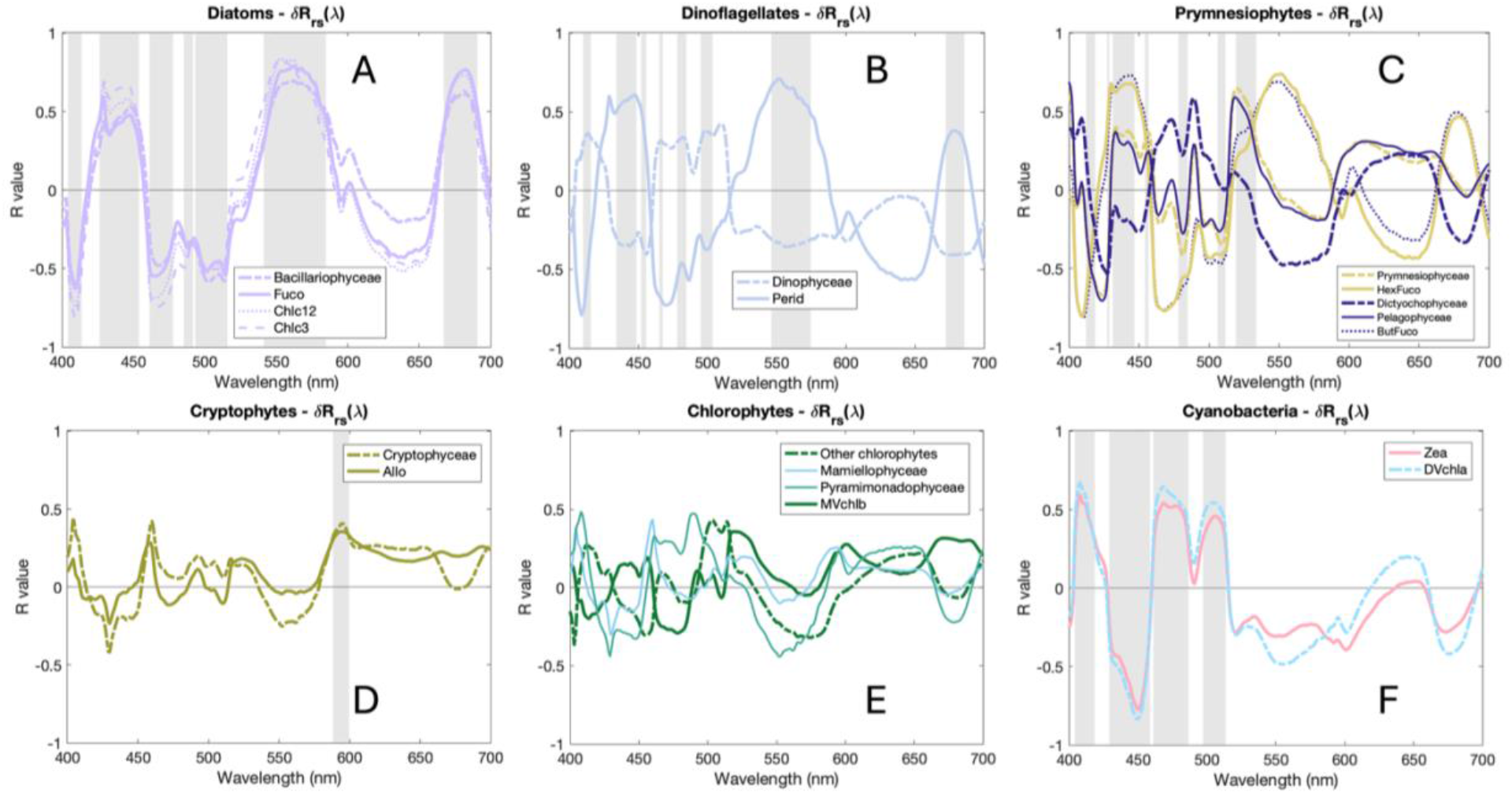
Spectral correlations in *δR_rs_*(*λ*) model coefficients for the major groups shared between 18S rRNA gene sequences and accessory pigments in this dataset: (A) diatoms, (B) dinoflagellates, (C) prymnesiophytes and silicoflagellates, (D) cryptophytes, (E) chlorophytes, (F) cyanobacteria (only identified by HPLC pigments, since bacteria do not have the 18S rRNA gene). Gray bars highlight wavelengths where the R value between the quantity (pigment or relative sequence abundance) in the model is significantly different from zero, indicating that this region is important for modeling.

## Conclusions

In this work, we tested whether hyperspectral ocean color data contain retrievable information about phytoplankton community composition by independently modeling pigments and 18S rRNA gene relative sequence abundances from in situ *R_rs_*(*λ*) data. While previous studies have retrieved broad phytoplankton groups based on pigment concentrations, this study directly modeled the eukaryotic phytoplankton community structure as assessed by metabarcoding. Despite the small dataset of paired global samples, the model retrievals for both pigments and genes were robust when compared to measured data. These results provide proof-of-concept that hyperspectral *R_rs_*(*λ*) contains information associated with eukaryotic phytoplankton community structure at the class level. The discrepancies between ocean color and PCC at some stations analyzed here (e.g., EXPORTS North Pacific) also reveal an important limitation: taxonomic variability cannot be recovered optically when it produces little corresponding change in the integrated optical signal.

Future studies should aim to increase the number of paired samples among the three methods assessed here (only possible with coordinated field campaigns and continued integration of public data), thereby expanding the range of ecosystems and phytoplankton community states represented and testing model transferability across regions and field campaigns, with the ultimate goal of developing global models for PACE and future hyperspectral ocean color missions. Additionally, next steps will include targeting a broader taxonomic range (i.e., including prokaryotic gene markers) either by combining methods (e.g., 18S rRNA gene sequences and 16S rRNA gene sequences; (Bei et al. 2025)) or using novel genetic markers and quantification protocols (e.g., *psbO* or JEDI; (Pierella Karlusich et al. 2022; McNichol et al. 2025)) or both. Together with expanding hyperspectral coverage provided by PACE, these observations will allow the challenges and opportunities of retrieving molecular PCC from space to be tested across much broader ecological gradients.

## Acknowledgments

SJK was supported by grants from NASA Ocean Biology and Biogeochemistry (grant numbers 80NSSC25K7428 and 80NSSC26K1128). REH was supported by the European Space Agency through the PlanktoSpace project (grant number 4000135756/21/I-EF), the French National Research Agency (ANR) through the Chaire de Professeur Junior program (ANR-22-CPJ1-0003-01), and the BNP Paribas Foundation through the PHYTOSCOPE project. Conversations with Emmanuel Boss, Colomban de Vargas, Ali Chase, Ivona Cetinić, and Colleen Durkin helped inform and improve this work. We are very grateful to everyone who worked to collect, analyze, and publicly share this data—global analyses are not possible without your help.

## Data availability statement

All data used in this work are publicly available. Tara Oceans HPLC pigments and remote sensing reflectance: https://seabass.gsfc.nasa.gov/experiment/Tara_Oceans_expedition. Tara 18S rRNA gene sequences: https://doi.org/10.5281/zenodo.7551643. NAAMES HPLC pigments,18S rRNA gene sequences, and remote sensing reflectance: https://seabass.gsfc.nasa.gov/experiment/NAAMES. EXPORTS HPLC pigments, 18S rRNA gene sequences, and remote sensing reflectance: https://seabass.gsfc.nasa.gov/experiment/EXPORTS. SDP model base code is available at: https://github.com/sashajane19/Rrs_pigments, https://github.com/max-danenhower/rrs-SDP-pigments, and https://nasa.github.io/oceandata-notebooks/notebooks/oci/oci_sdp.html.

## Supplemental Figures

**Figure S1.**
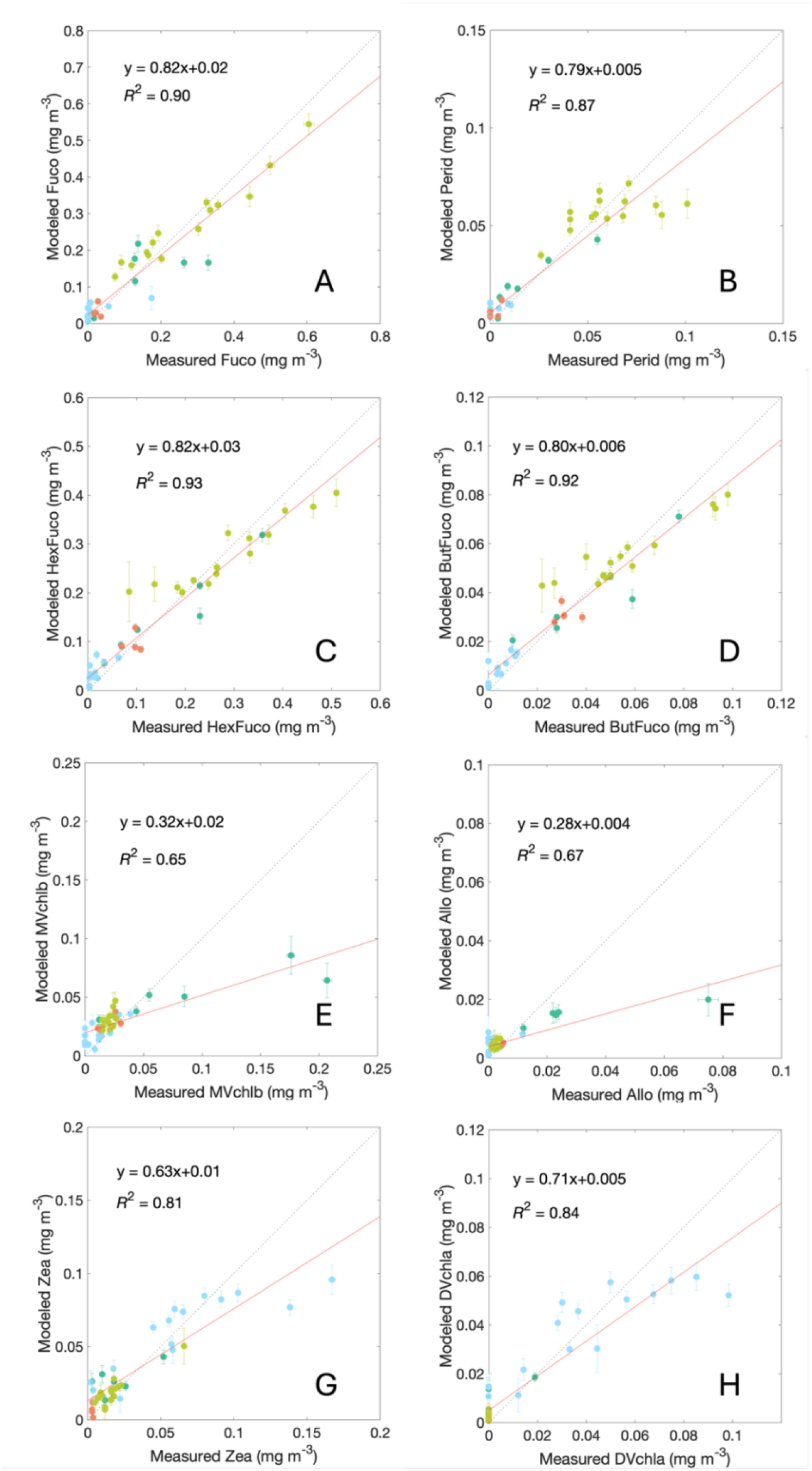
Relationship between measured and modeled pigments for (A) Fuco, (B) Perid, (C) HexFuco, (D) ButFuco, (E) MVchlb, (F) Allo, (G) Zea, (H) DVchla. Measured error bars show the range of measurement certainty (Van Heukelem and Thomas 2001) and modeled error bars show the 95% CI of the 100 repeated train-validation partitions.

**Figure S2.**
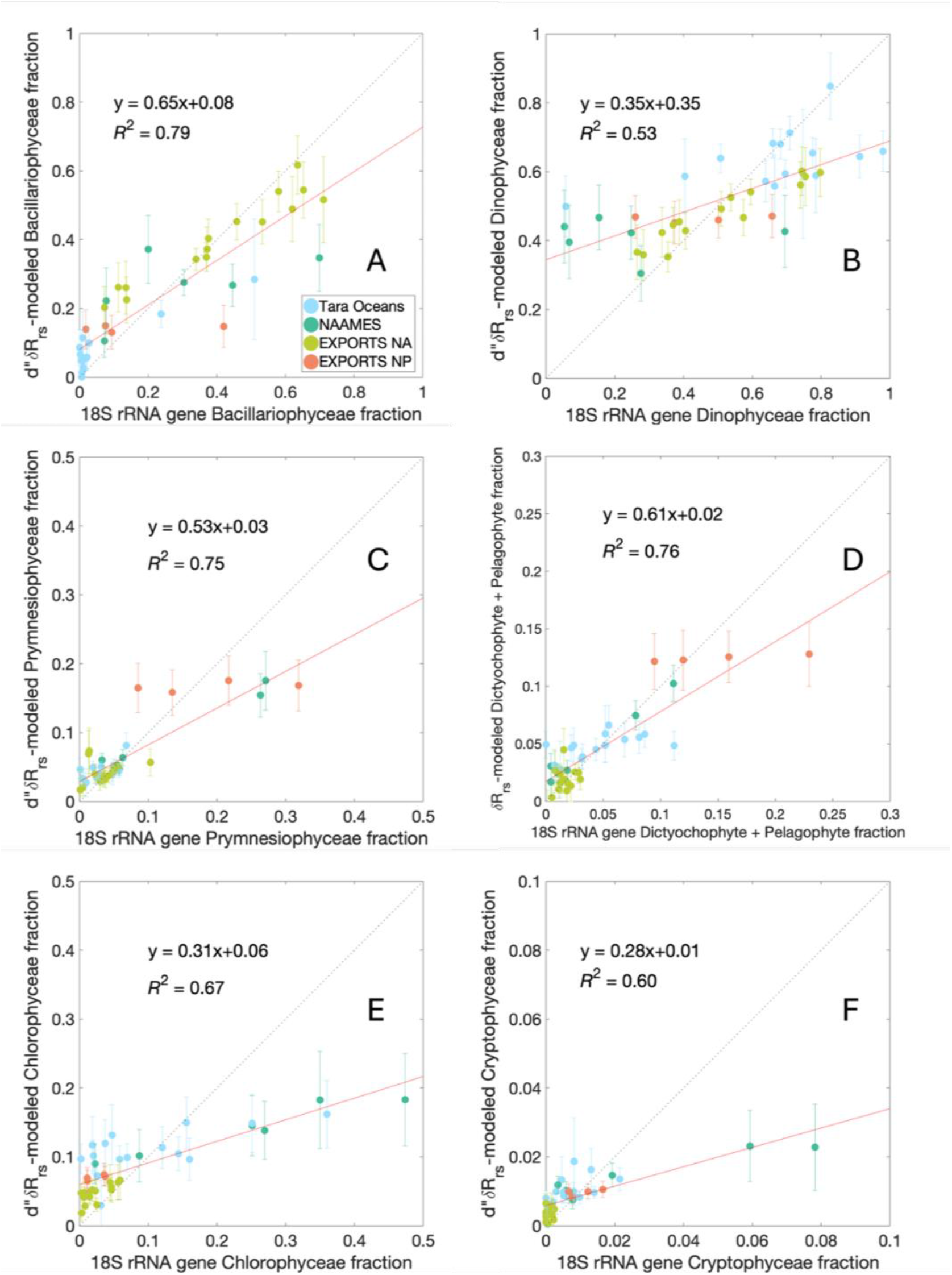
Relationship between measured and modeled relative 18S rRNA gene class abundances for (A) diatoms, (B) dinoflagellates, (C) prymnesiophytes, (D) dictyochophytes + pelagophytes, (E) chlorophytes, and (F) cryptophytes. Modeled error bars show the 95% CI of the 100 repeated train-validation partitions.

**Figure S3.**
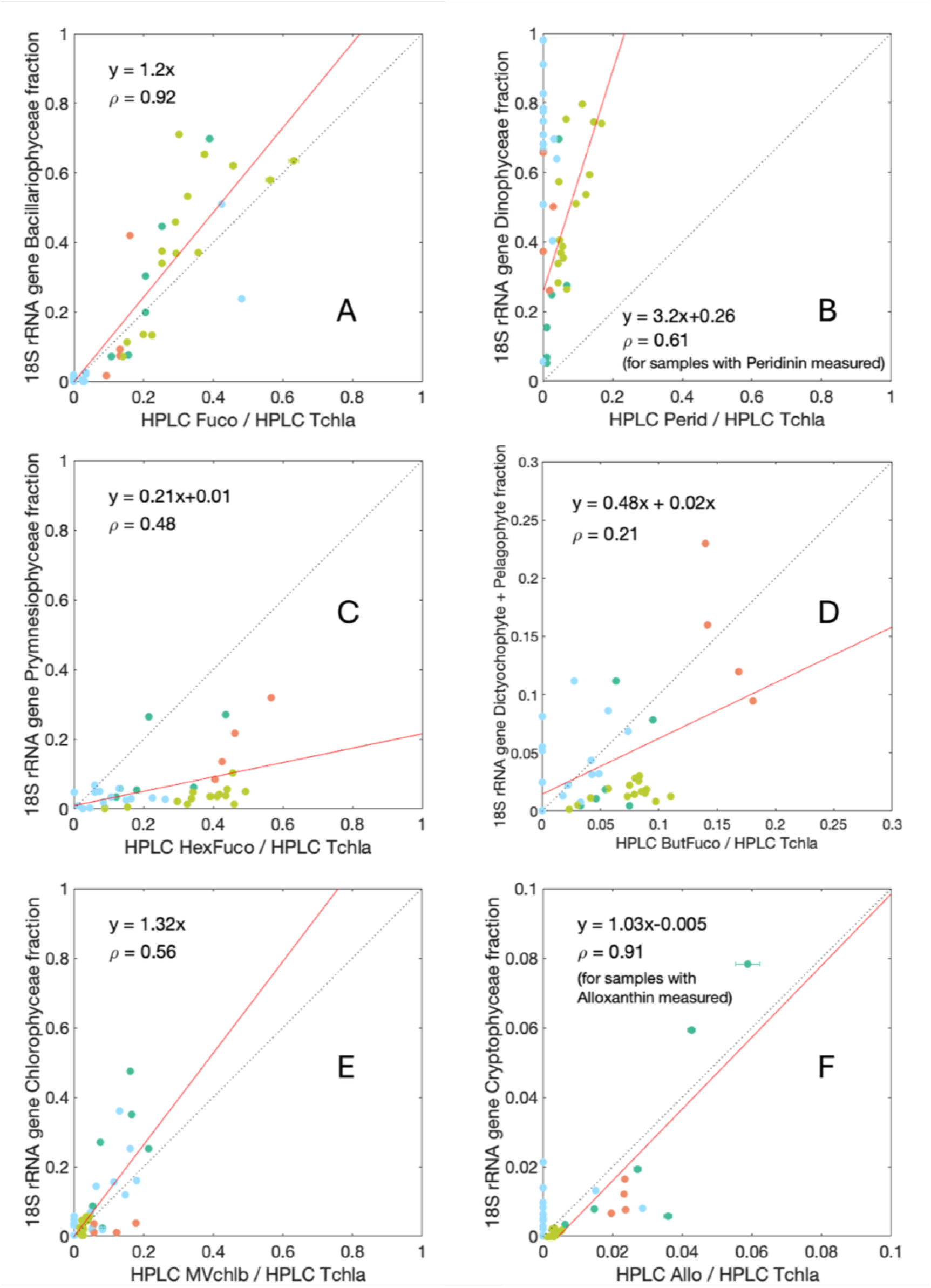
Relationship between HPLC pigments and relative 18S rRNA gene sequence abundances for the groups shared between pigments and genes: (A) diatoms, (B) dinoflagellates, (C) prymnesiophytes, (D) dictyochophytes + pelagophytes, (E) chlorophytes, and (F) cryptophytes.

